# Genome analysis leads to discovery of Quorum Sensing Genes in *Cedecea neteri*

**DOI:** 10.1101/018887

**Authors:** Kian-Hin Tan, Kok-Gan Chan

## Abstract

We have identified a strain of C. neteri SSMD04 isolated from pickled mackerel sashimi that produced N-acyl-homoserine lactone (AHL) type quorum sensing (QS) activity. Tandem mass sspectrometry revealed that C. neteri SSMD04 produced N-butyryl-homoserine lactone (C4-HSL). We identified a pair of luxIR homologues in this genome that shares the highest similarity with croIR from Citrobacter rodentium. The AHL synthase, which we named it as cneI (636 bp) and at 8bp distance from cneI is a sequence encoding a hypothetical protein, potentially the cognate receptor, a luxR homologue which we named it as cneR. We also found an orphan luxR in this isolate. To our knowledge, this is the first report on the AHL production activity in C. neteri, discovery of its luxI/R homologues and the orphan receptor.

## 1. Introduction

*Cedecea* spp. are extremely rare Gram-negative bacteria that belong to the *Enterobacteriaceae* family (Berman, 2012). This genus is lipase-positive and resistant to colistin and cephalothin. The name *Cedecea* was coined by P. A. D. Grimont and F. Grimont, from the abbreviation of the Centers for Disease Control (CDC) (Grimont, Grimont & Farmer, 1981). Originally recognized as Enteric group 15, this genus is comprised of five species, out of which only three were named, *C. neteri*, *C.lapagei*, *C. davisae*, and the other two are known as *Cedecea* species 3 and *Cedecea* species 5 (Brenner et al., 2005).

*Cedecea* species 4 was given the name *C. neteri* in 1982 when its clinical significance was reported. The name ‘*neteri*’ was coined to honor Erwin Neter, an American physician-microbiologist for his contributions in the work on *Enterobacteriaceae* in human disease (Farmer III et al., 1982). *C. neteri* was also found in a patient with systemic lupus erythematosus where it led to the patient’s death (Aguilera et al., 1995). Even though it was evident that *C. neteri* can act as human pathogen, its etiology is unknown and limited studies have been conducted on *Cedecea* spp. There were cases of isolation of *Cedecea* spp. from other sources except human (Jang & Nishijima, 1990; Osterblad et al., 1999), and we have recently reported the isolation of *C. neteri* SSMD04 from *Shime saba* (Chan et al., 2014), a Japanese cuisine that involves marinating with salt and rice vinegar, enabling the usually perishable *saba* (mackerel) to be enjoyed in the form of *Sashimi* (raw fish).

Bacteria demonstrate a concerted gene regulation mechanism termed ‘Quorum Sensing’ (QS) that relies on the population density of the bacteria (Fuqua, Winans & Greenberg, 1996; Miller & Bassler, 2001; Schauder & Bassler, 2001). The mechanism of QS involves the production, release, detection, and response to small diffusible molecules known as autoinducers, such as *N*-acyl homoserine lactones (AHLs) commonly employed by Gram negative bacteria (Chhabra et al., 2005; Williams et al., 2007). AHL molecules are generally characterized by the length and saturation of its acyl side chains which can vary from 4 to 18 carbons (Pearson, Van Delden & Iglewski, 1999), as well as the R-group substitution at the third carbon (Pearson, Van Delden & Iglewski, 1999; Waters & Bassler, 2005). QS has been shown to play a role in the regulation of a wide range of phenotypes, such as antibiotic biosynthesis, biofilm formation, pathogenesis, bioluminescence, antibiotic production and more (Fuqua, Winans & Greenberg, 1996; Salmond et al., 1995; de Kievit & Iglewski, 2000, Hastings & Nealson, 1977; Bainton et al., 1992; Eberl et al., 1996).

## 2. Materials and methods

### 2.1 Sample collection and processing

*Shime saba sashimi* sample was collected from a local supermarket in Malaysia and processed within half an hour following collection. Five grams of sample was stomached and homogenized in 50 ml of peptone water and then spread on MacConkey (MAC) agar. The culture plates were incubated overnight in 28 °C.

### 2.2 Bacterial strains, media and culture conditions

*C. neteri* SSMD04, *Chromobacterium violaceum* CV026, *Erwinia carotovora* GS101 and *E. carotovora* PNP22 were maintained in Luria Bertani (LB) medium at 28 °C. *lux*-based biosensor *Escherichia coli* [pSB401] was grown in LB supplemented with tetracycline (20 μg/mL) at 37 °C. All broth cultures were incubated with shaking (220 rpm).

### 2.3 Species identification of isolate SSMD04

#### 2.3.1 16S rDNA phylogenetic analysis

16S rDNA sequence was extracted from the complete genome sequence of isolate SSMD04, while other 16S rDNA sequences of *Cedecea*. spp. were retrieved from GenBank. The Molecular Evolutionary Genetics Analysis (MEGA) 6.0 (Tamura et al., 2013) was used to align the sequences and construct a Maximum likelihood tree using 1,000 bootstrap replications.

#### 2.3.2 Biolog GEN III microbial identification system

Microbial identification using Biolog GEN III MicroPlate^TM^ was carried out according to manufacturer’s protocol. In brief, overnight culture of *C. neteri* SSMD04 grown on Tryptic Soy Agar (TSA) was used to inoculate inoculating fluid (IF) A to a cell density of 90-98% transmittance. The inoculum was then pipetted into each well of the MicroPlate^TM^ (100 μL per well) and incubated at 28 °C for 24 hrs. The MicroPlate was then read using Biolog’s Microbial Identification Systems software where the wells will be scored as ‘negative’ or ‘positive’ based on the colour change due to the reduction of tetrazolium redox dyes. This ‘Phenotypic Fingerprint’ was then used to identify the bacteria by matching it against the database in the system.

### 2.4 Detection of AHL production in *C. neteri* SSMD04

AHL-type QS activity of *C. neteri* SSMD04 was screened using biosensor *C. violaceum* CV026. This is performed by cross streaking *C. neteri* SSMD04 against *C. violaceum* CV026. *E. carotovora* GS101 and *E. carotovora* PNP22 were used as positive and negative controls, respestively (McClean et al., 1997).

### 2.5 AHL extraction

*C. neteri* SSMD04 was cultured overnight at 28 °C in LB broth (100 mL) supplemented with 50 mM of 3-(N-morpholino)propanesulfonic acid (MOPS) (pH5.5). Spent supernatant was collected by centrifugation and subsequently extracted twice with equal volume of acidified ethyl acetate (AEA) (0.1 % v/v glacial acetic acid). The extracts were air dried and reconstituted in 1 mL of AEA, transferred into sterile microcentrifuge tubes and air dried again, before being stored at -20 °C. The extracts were later used for detection of AHL by *lux*-based biosensor *E. coli* [pSB401] as well as triple quadrupole LC/MS.

### 2.6 AHL identification by triple quadrupole LC/MS

Extracts from section 2.5 were reconstituted in acetonitrile (ACN) prior to LC/MS analysis as described before (Lau et al., 2013) with slight modification. In brief, mobile phase A used was water with 0.1 % v/v formic acid and mobile phase B used was ACN with 0.1 % formic acid. The flow rate used was 0.5 mL/min. The gradient profile was set to: A:B 80:20 at 0 min, 50:50 at 7 min, 50:50 at 7.10 min, 80:20 at 12 min, 80:20 at 12.10 min, 20:80 at 14 min, 20:80 at 14.10 min. Precursor ion scan mode was carried out in positive ion mode with Q1 set to monitor *m/z* 90 to *m/z* 400 and Q3 set to monitor for *m/z* 102. ACN was also used as a blank.

### 2.7 Measurement of bioluminescence

*E. coli* [pSB401] (Winson et al., 1998) was used as biosensor for the detection of exogenous short chain AHLs present in the extracts. The biosensor strain was cultured in LB broth supplemented with tetracycline (20 μg/mL). The overnight culture was then diluted to an OD_600_ of 0.1 with fresh LB broth with tetracycline. The diluted *E. coli* culture was used to resuspend the extracts from section 2.5, prior to being dispensed into a 96-well optical bottom microtitre plate. Cell density bioluminescence measurements were carried out by Infinite M200 luminometer-spectrophotometer (Tecan, Männedorf, Switzerland) over a period of 24 hours. Diluted *E. coli* culture without extracts was read for normalization and sterile broth was used as negative control. The results were displayed as relative light units (RLU)/OD495 nm against incubation time.

### 2.8 Genome annotation and analysis

Whole genome of *C. neteri* SSMD04 was annotated by RAST as described (Chan et al., 2014). DNAPlotter (Carver et al., 2009) was used to construct GC plot and GC skew.

## 3. Results

### 3.1 Species identification of *C. neteri* SSMD04

16S rDNA sequence retrieved from whole genome sequence of *C. neteri* SSMD04 was used to construct a phylogenetic tree with other sequences of *Cedecea* spp. available in GenBank. 16S rDNA sequence of *C. neteri* SSMD04 clusters with other *C. neteri* strains in a monophyletic clade (Figure 1). However, it can also be observed that the available 16S rDNA sequences of *C. davisae* formed two distinct clusters, of which one is more closely related to *C. neteri* and the other *C. lapagei*.

**Figure 1.**
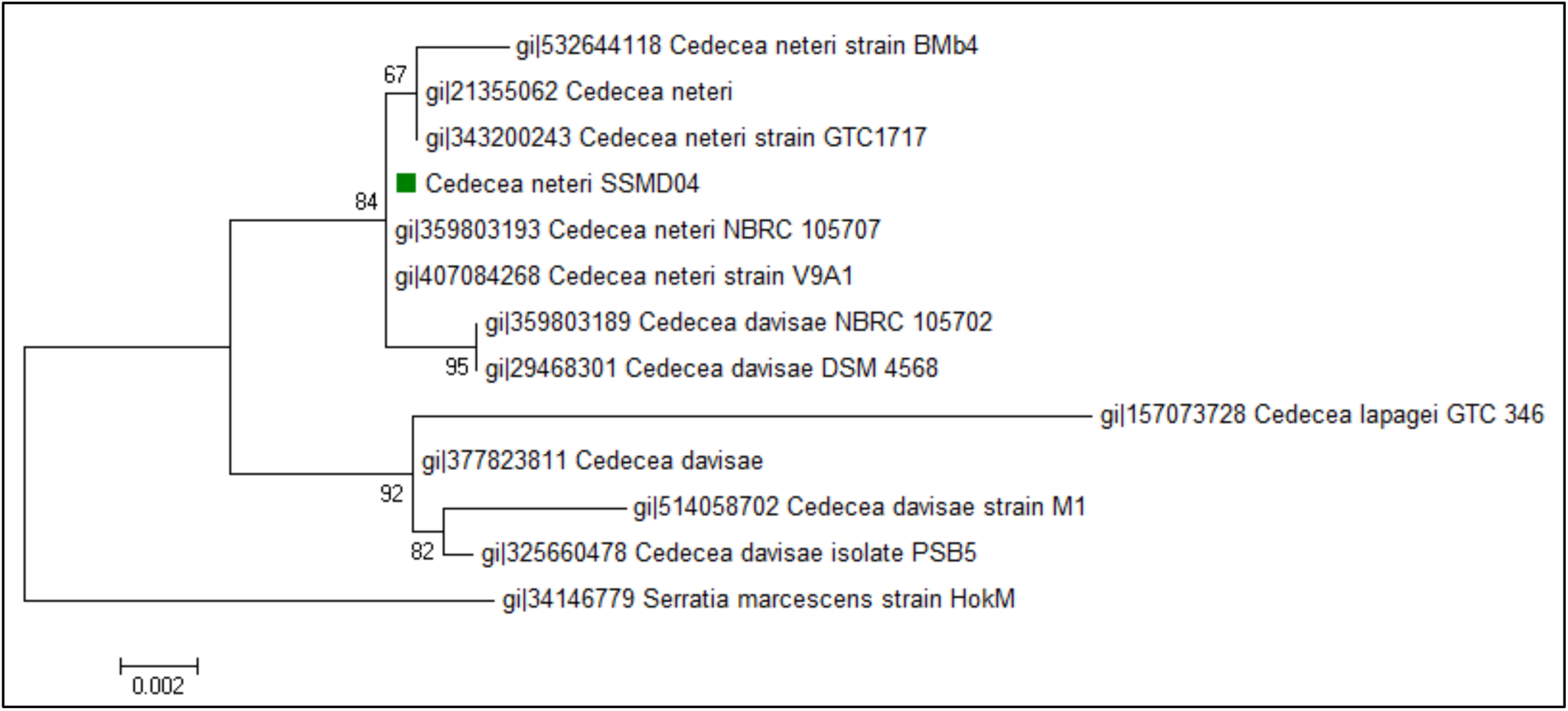
Phylogenetic tree showing the position of *C. neteri* SSMD04 (green square) relative to other *Cedecea* spp. The neighbour joining tree was inferred from 1,297 aligned positions of the 16S rDNA sequences using Hasegawa-Kishino-Yano substitution model. Boostrap values are represented at the branch point. The scale denotes the number of substitutions per nucleotide position. *Serratia marcescens* strain HokM was used as an outgroup.

Biology Gen III microbial identification system was also used to assess the identity of *C. neteri* SSMD04 biochemically. The system identified this strain to be *C. neteri* with a probability and similarity of 0.697. The positive reaction in sucrose well and D-sorbitol well agrees with the report of Farmer III et al., 1982.

### 3.2 Detection of AHL-type QS activity in *C. neteri* SSMD04 using AHL biosensor

*C. neteri* SSMD04 was streaked on LBA against biosensor *C. violaceum* CV026. Short chain AHLs produced from *C. neteri* SSMD04 diffused passively towards the biosensor, activating the production of purple colour pigment, violacein. The result shown in Figure 2 indicated the presence of exogenous AHLs in *C. neteri* SSMD04 culture.

**Figure 2.**
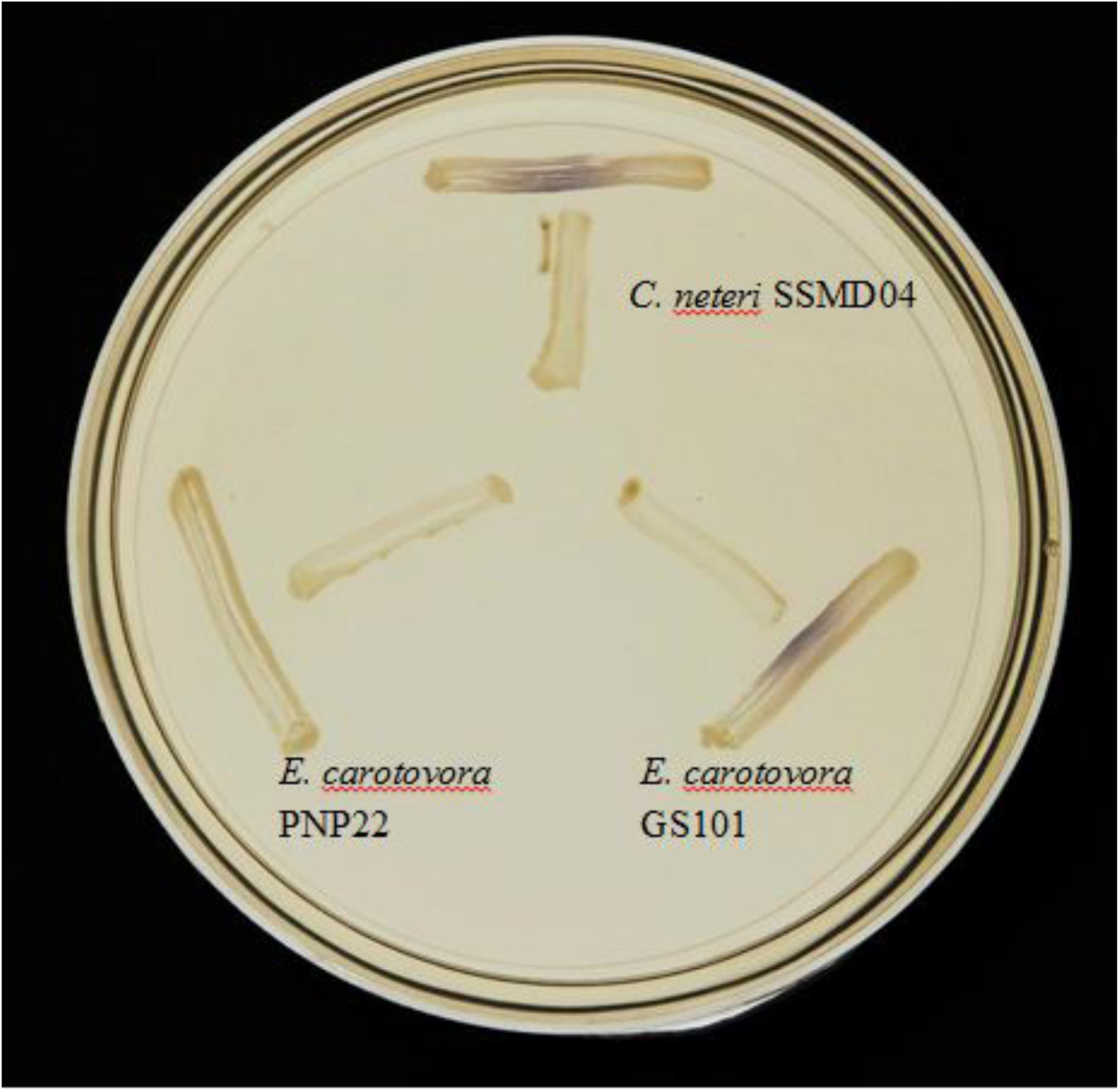
Screening for AHL-type QS activity of *C. neteri* SSMD04 using the biosensor *C. violaceum* CV026. *E. carotovora* PNP22 and *E. carotovora* GS101 act as negative and positive controls, respectively.

### 3.3 Measurement of bioluminescence

*C. neteri* SSMD04 also activated *lux*-based biosensor *E. coli* [pSB401] which produces bioluminescence in the presence of short chain AHLs (Figure 3).

**Figure 3.**
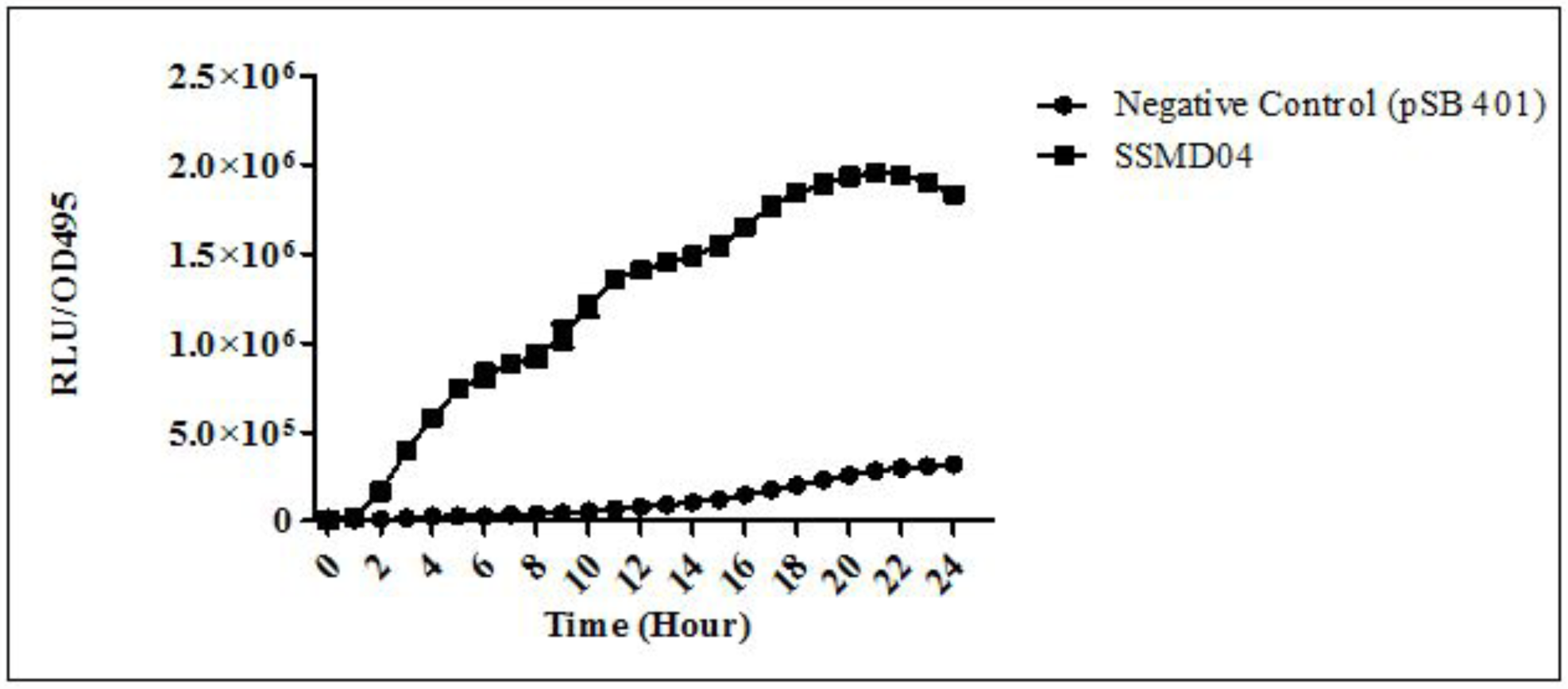
Relative light unit (RLU)/OD495 against incubation time of cultures of *E. coli* [pSB401] in the presence of extracted AHLs (square plots) and negative control (circle plots).

### 3.4 AHL identification by triple quadrupole LC/MS

The extracted-ion chromatogram (EIC) generated from the triple quadrupole LC/MS system showed a peak with the same retention time as that of the synthetic *N*-butyryl-homoserine lactone (C4-HSL) standard, which was constantly present in all three replicates (Figure 4). Analysis of the mass spectrum (MS) data revealed the presence of a peak with mass-to-charge ratio (*m/z*) of 172 (Figure 5), which is consistent with the previously reported value (Ortori et al., 2011). This implication was strengthened by the presence of a product ion peak (*m/z* = 102).

**Figure 4.**
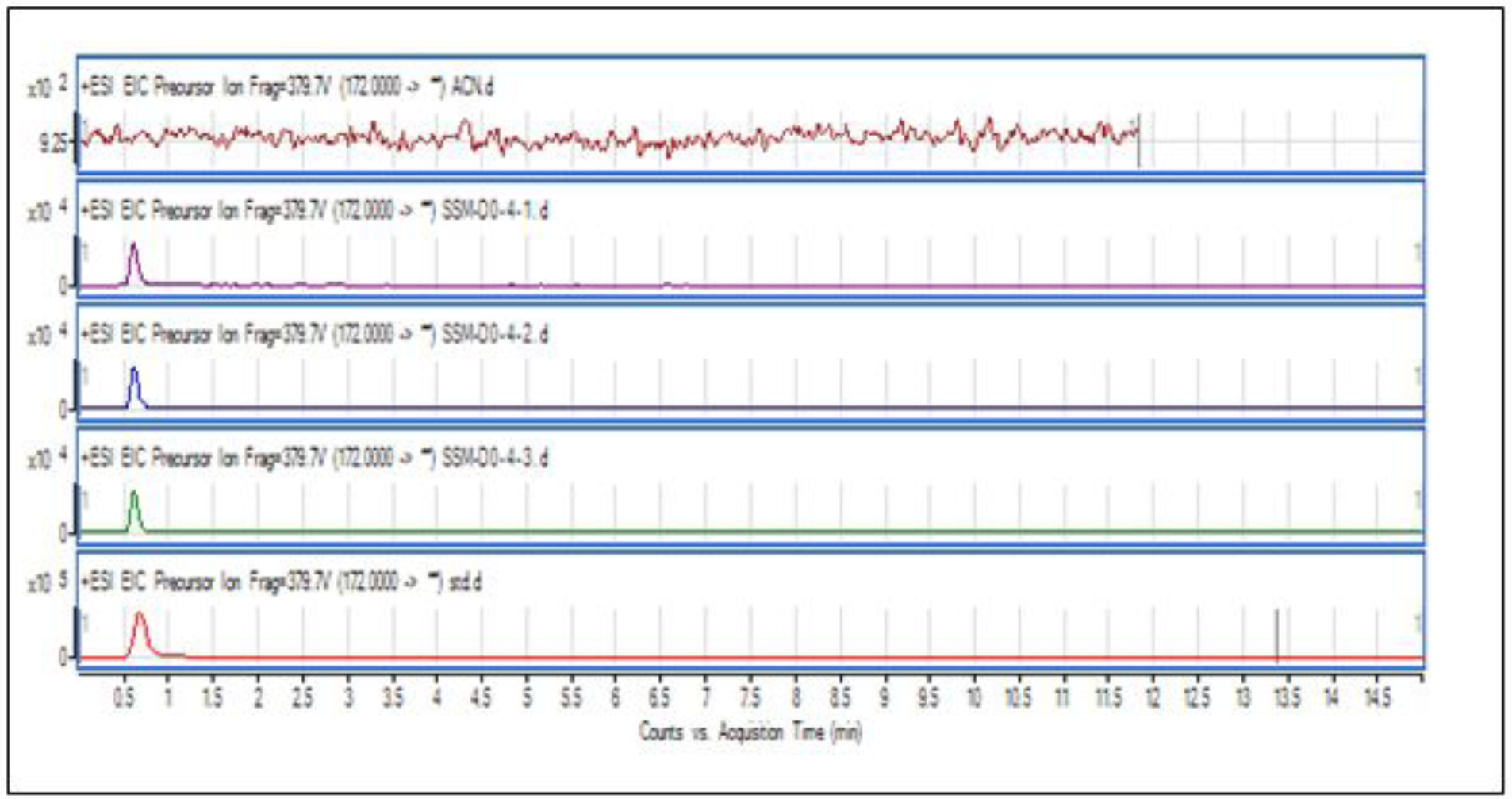
EIC of *C. neteri* SSMD04 extract. The data represented three replicates of the extract against synthetic C4-HSL. ACN was used as blank.

**Figure 5.**
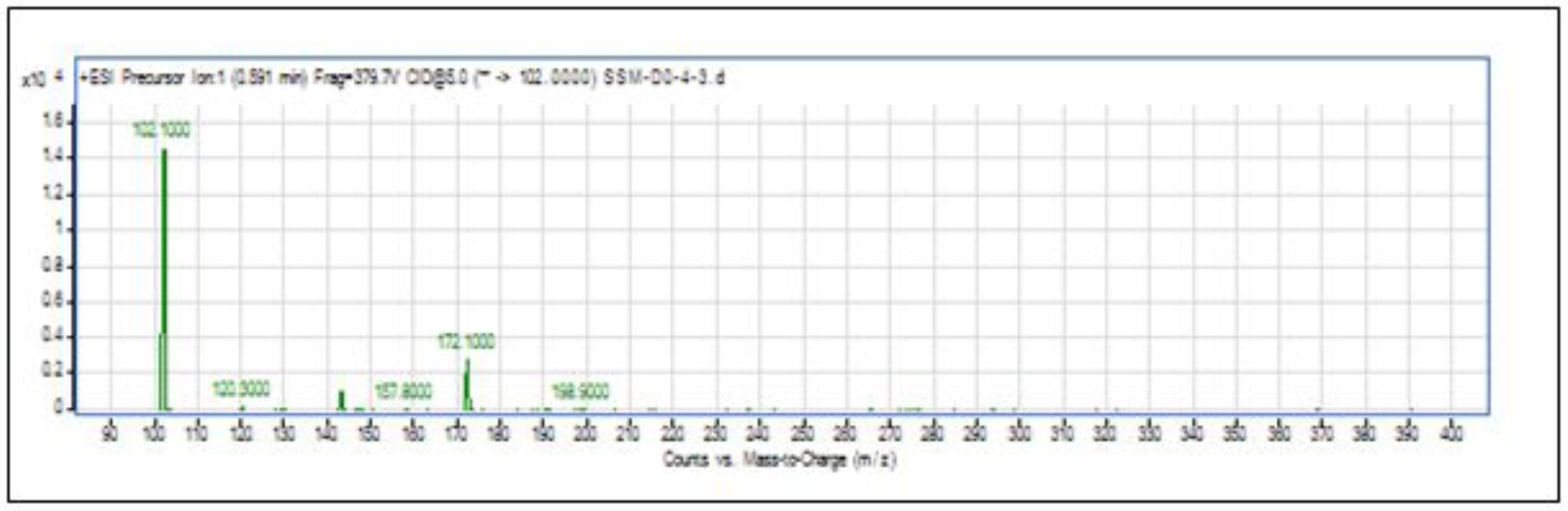
Product ions of the peak seen in Figure 4. This shows that the extract of *C. neteri* SSMD04 contains C4-HSL.

### 3.5 Genome annotation and analysis

As reported previously, the whole genome sequence of *C. neteri* SSMD04 was annotated by RAST (Aziz et al., 2012) (Figure 6). Figure 7 shows the visualization of *C. neteri* SSMD04 genome. From the data generated by NCBI prokaryotic genome annotation pipeline, a 636 bp *luxI* homologue, hereafter named *cneI*, was found in the genome. This gene shares 70 % base pair similarity with *N*-acyl homoserine lactone synthase *croI* of *Citrobacter rodentium* ICC168. Analysis of amino acid sequence of *cneI* using InterPro (Mitchell et al., 2015) identified the presence of an acyl-CoA-*N*-acyltransferase, the structural domain of *N*-acyl homoserine lactone synthetases (Gould, Schweizer and Churchill, 2004; Watson et al., 2002).

**Figure 6.**
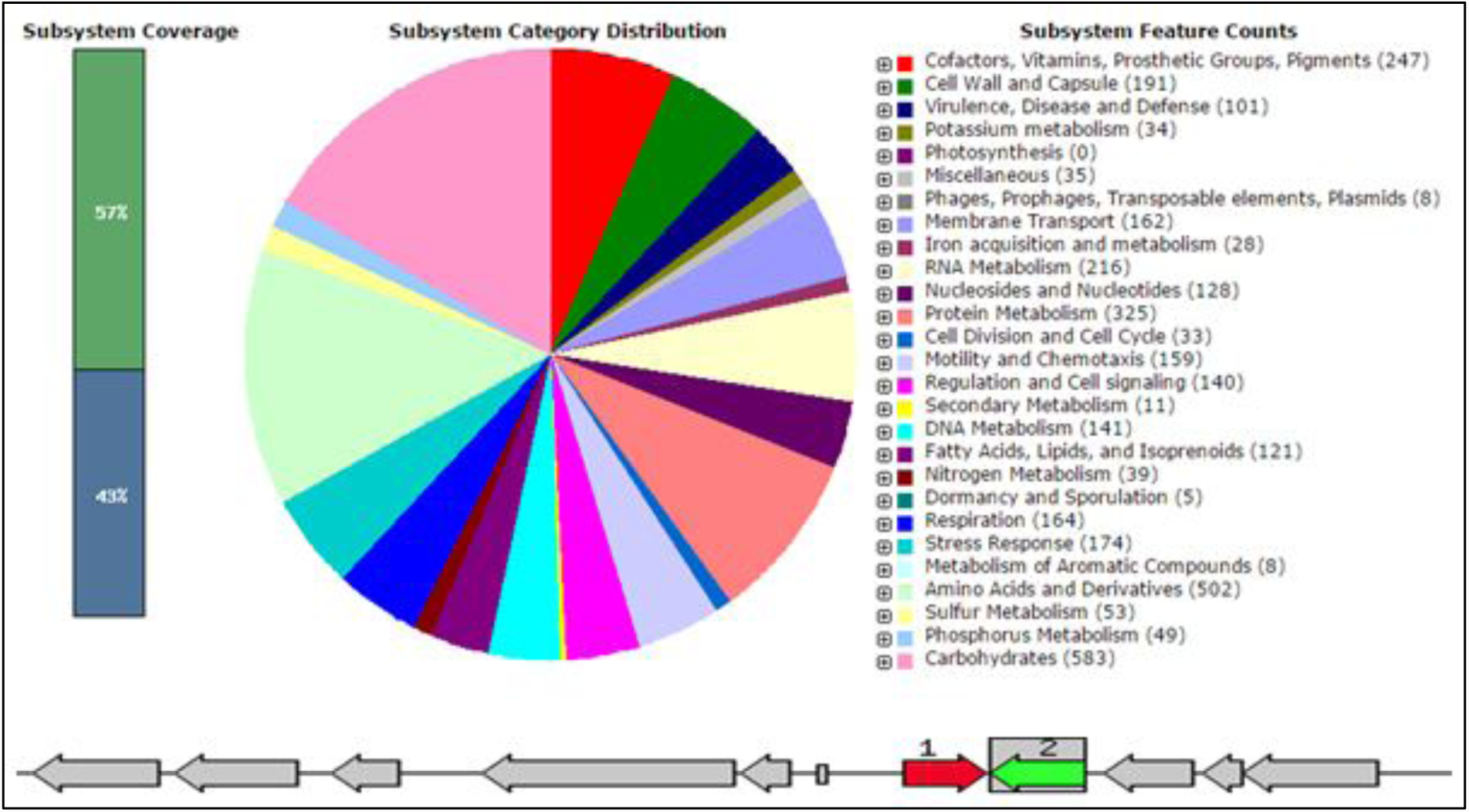
From left to right: Alkanesulfonate utilization operon LysR-family regulator Cbl; Nitrogen assimilation regulatory protein Nac; Membrane protein, suppressor for copper-sensitivity ScsD; Membrane protein, suppressor for copper-sensitivity ScsB; Suppression of copper sensitivity: putative copper binding protein ScsA; tRNA-Asn-GTT; LuxI homologue protein; LuxR homologue protein; Oxygen-insensitive NAD(P)H nitroreductase/ Dihydropteridine reductase; hypothetical protein; Bifunctional protein: zinc-containing alcohol dehydrogenase/ quinone oxidoreducatse (NADPH: quinone reductase).

Adjacent to *cneI*, 8 bp away, is a sequence encoding a hypothetical protein, potentially the cognate receptor, a *luxR* homologue (*cneR*). The coding region was found to be 705 bp long and share 70 % similarity with *croR* of *C. rodentium*. Analysis of this protein reveals two signature domains, the autoinducer-binding domain and the C-terminal effector.

**Figure 7.**
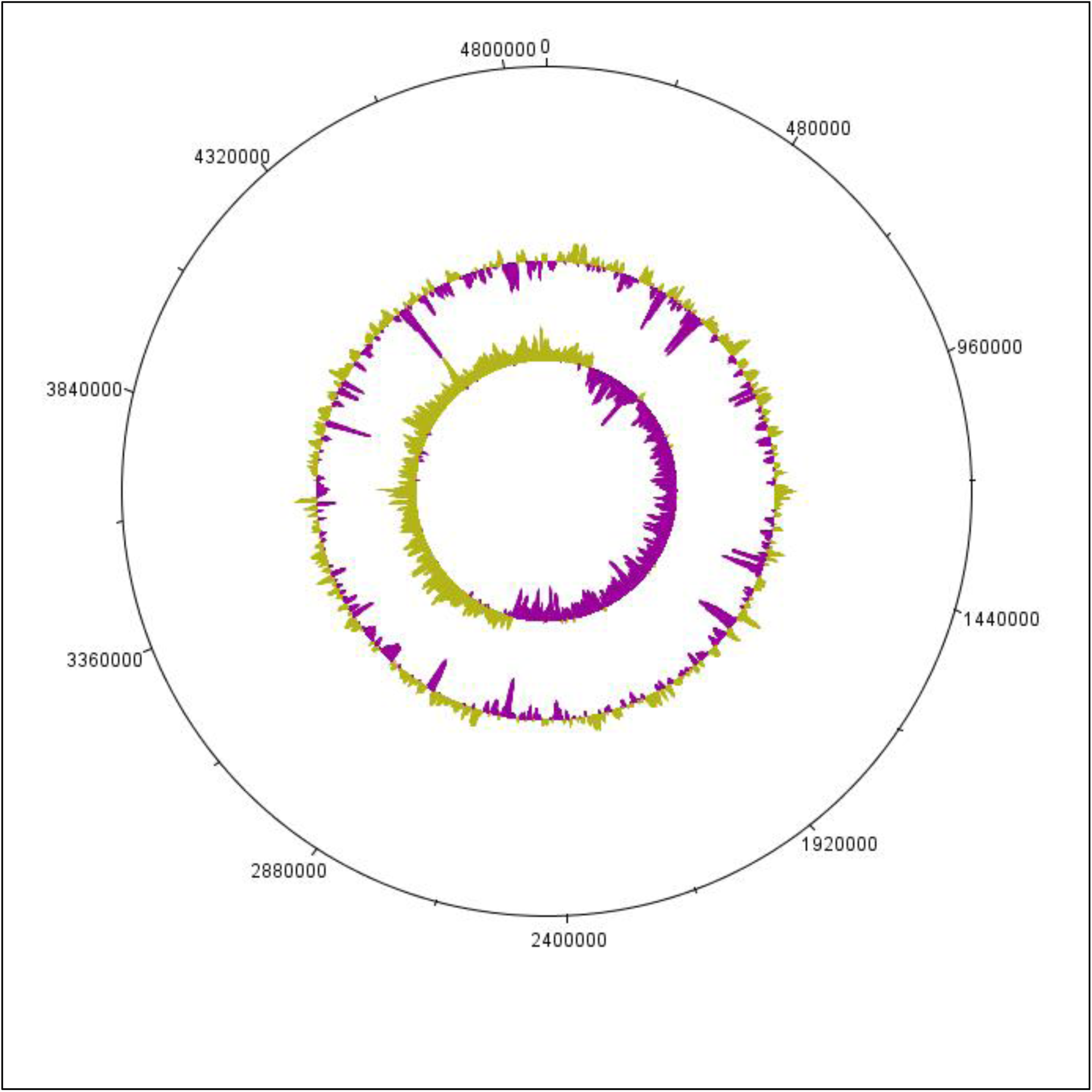
Whole genome representation of *C. neteri* SSMD04 genome using DNAplotter. The number on the outermost circle is the scale for the plot. The innermost circle represents GC skew, while the second circle represents GC content.

Apart from that, a sequence potentially encoding an orphan *luxR* type receptor (723 bp) was also found within the genome. This luxR homologue shares 69% sequence homology to *luxR* homologue of *Enterobacter asburiae* L1.

## 4. Discussion

*Cedecea* spp. are very rare bacteria and thus not well studied. Despite the evidence of their ability to infect human, their medical significance can be overlooked due the poor understanding of their physiology and etiology. Besides that, they are potentially challenging pathogens due to their resistance towards a wide range of antimicrobial agents, such as cephalothin, extended spectrum cephalosporins, colistin, and aminoglycosides (Mawardi et al., 2010; Abate, Qureshi & Mazumder, 2011; Dalamaga et al., 2008). To date, isolation of this species from non-clinical source has not been reported. Therefore, the isolation of *C. neteri* SSMD04 from a food source expands the current knowledge on diversity of the genus *Cedecea*.

Although employed by a wide range of Gram-negative bacteria in gene regulation that allows the alteration of behaviour on a population level (Waters & Basslers, 2005), AHL-type QS activities have not been reported in *C. neteri*. Some bacteria utilizes QS to regulate virulence and thus gaining advantage of expressing virulence factors only when the population density is large enough to triumph the hosts’ immune system (Passador et al., 1993; Brint & Ohman, 1995; McClean et al., 1997; Thomson et al., 2000; Weeks et al., 2010). Given the understanding that *C. neteri* can act as human pathogen, it can be hypothesized that AHL-type QS activity in *C. neteri* is involved in the regulation of virulence factors. However, further studies on clinical as well as non-clinical strains would help in the solution of this hypothesis.

The whole genome sequence provides very valuable information in studying the genetic basis of QS in *C. neteri* SSMD04. The finding of *cneIR* in this genome, lying adjacent to each other, demonstrated a common feature of *luxIR* homologues (Brint & Ohman, 1995; Fuqua, Winans & Greenberg, 1994; Williamson et al., 2005). Analysis of amino acid sequence of the *cneIR* pair with InterPro agreed with their identity. The *cneIR* pair was found to be most similar to *croIR* in *C.rodentium*. *C. rodentium* has been found to produce C4-HSL as the major AHL and C6-HSL as the minor (Coulthurst et al., 2007). *C. neteri* SSMD04 also produces C4-HSL, but it does not produce detectable level of C6.

The presence of lipase-positive *C. neteri* in marinated oily fish strongly suggests its role as a potential food spoilage agent, not only because of its ability to survive an extreme environment of high salinity and acidity, but also the fact that AHLs have long been associated with food spoilage via regulation of the proteolytic and lipolytic pathways (Skandamis & Nychas, 2012; Bruhn et al., 2004). Nevertheless, the roles of *C. neteri* in pathogenesis and food spoilage still require more information to be elucidated.

## 5. Conclusion

This study has confirmed the production of C4-HSL by *C. neteri* SSMD04 isolated from *Shime saba sashimi*. This is the first report of QS activity in *C. neteri*. However, the function of QS in *C. neteri* SSMD04 is still unknown. We hope that further studies coupled with the available genome information of *C. neteri* SSMD04 can help to elucidate the regulatory circuit of *C. neteri* SSMD04 by QS.

